# PLAbDab-nano: a database of camelid and shark nanobodies from patents and literature

**DOI:** 10.1101/2024.07.19.604232

**Authors:** Gemma L. Gordon, Alexander Greenshields-Watson, Parth Agarwal, Ashley Wong, Fergus Boyles, Alissa Hummer, Ana G. Lujan Hernandez, Charlotte M. Deane

## Abstract

Nanobodies are essential proteins of the adaptive immune systems of camelid and shark species, complementing conventional antibodies. Properties such as their relatively small size, solubility and high thermostability make VHH and VNAR modalities a promising therapeutic format and a valuable resource for a wide range of biological applications. The volume of academic literature and patents related to nanobodies has risen significantly over the past decade. Here, we present PLAbDab-nano, a nanobody complement to the Patent and Literature Antibody Database (PLAbDab). PLAbDab-nano is a self-updating, searchable repository containing approximately 5000 annotated VHH and VNAR sequences. We describe the methods used to curate the entries in PLAbDab-nano, and highlight how PLAbDab-nano could be used to design diverse libraries, as well as find sequences similar to known patented or therapeutic entries. PLAbDab-nano is freely available as a searchable web server (*opig*.*stats*.*ox*.*ac*.*uk/webapps/plabdab-nano/*).

**Graphical Abstract:** 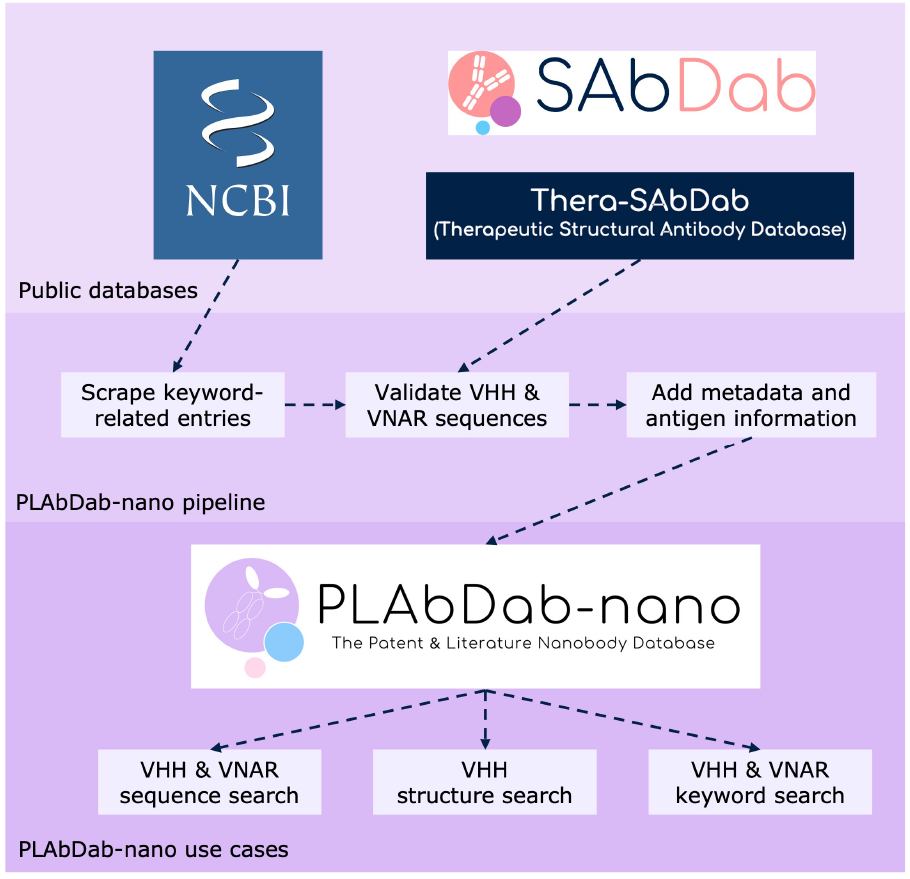

## Introduction

Owing to advantageous properties such as their relatively small size, high solubility and thermostability (*1–3*), nanobodies have increasingly garnered interest as a potential therapeutic format. Although, to date, only a handful of nanobody-based therapies have been approved, there are many progressing through the stages of clinical development (*4, 5*). In addition to their use as drugs, nanobodies are valuable tools in other areas of medical and scientific research, for example, as crystallization chaperones, in diagnostics, and for imaging (*4, 6*).

Nanobodies are derived from the antigen-binding portion of heavy-chain antibodies (HCAbs), which are produced by the adaptive immune systems of camelid and shark species. There are two varieties of nanobody: the VHH (the Variable Heavy domain of the Heavy chain) which is derived from camelids, and the VNAR (Variable New Antigen Receptor), which is derived from sharks (**Figure 1**). VHH nanobodies share a closer evolutionary lineage to the heavy chain of conventional IgG antibodies than VNAR nanobodies.

**Figure 1.**
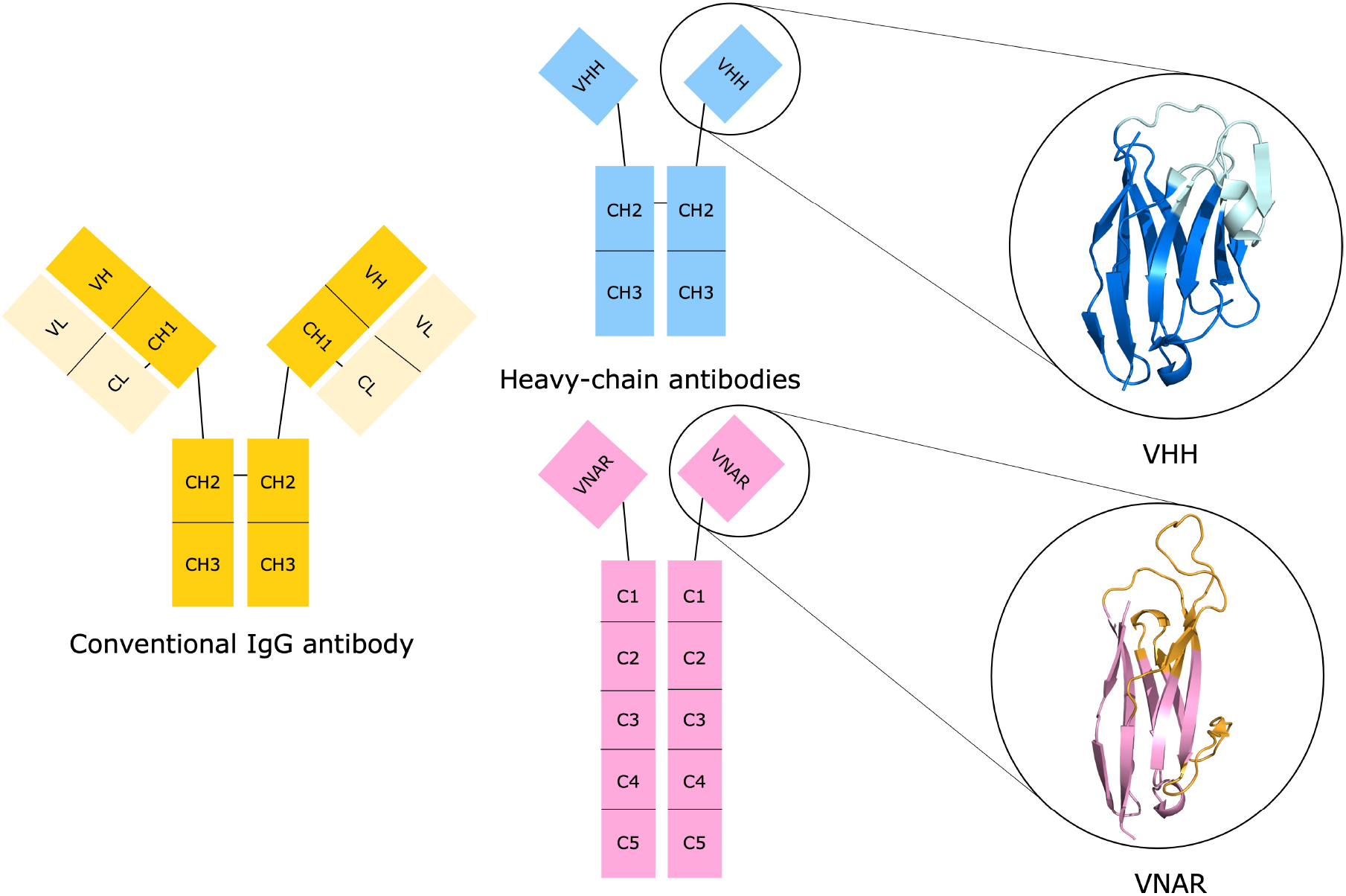
Nanobodies are derived from the antigen-binding portion of heavy-chain antibodies (HCAbs), which lack the light chain pairing possessed by conventional antibodies (shown in yellow). There are two varieties of nanobody: the VHH from camelid species (shown in blue, with the lighter blue denoting the CDR loops), and the VNAR from sharks (shown in pink, with the CDR loops and hypervariable regions shown in orange). PDB entries 8HR2 and 7S83 were used to create the VHH and VNAR figures.

Within the VHH structure, the binding site (paratope) is concentrated primarily in three hypervariable loops, known as the complementarity-determining regions (CDRs). The CDR3, which is longer in nanobodies compared to conventional antibodies (*7–12*), contributes most to the binding site (*12*). Framework residues also play a significant role in nanobody binding, being incorporated into the paratope more frequently than is observed for conventional antibodies (*12–14*). VNARs possess a structurally analogous scaffold to VHHs, but differ in that they feature only two CDR loops (CDR1 and CDR3), alongside two additional hypervariable regions (HV2 and HV4) (*15, 16*). Previous work has sought to collect nanobody data from many sources. For example, the antibody sequence database OAS (*17, 18*) aims to collect immune repertoire data and contains 1.6 million VHH sequences, but as these are from repertoire studies, they do not have any functional annotation. The structural data for 1493 unique VHH sequences is available in SAbDab-nano (*19*). TheraSAbDab is a collection of therapeutic antibodies, and contains 30 VHH or single-domain antibody entries (*20*). CoV-AbDab, a database for antibodies targeting the COVID-19 virus, contains 801 nanobodies (*21*).

Other repositories dedicated solely to nanobodies have also been curated, drawing from multiple public sources, such as sdAb-DB (*22*), INDI (*23*), and most recently, NanoLAS (*24*). SdAb-DB contains only a small number of entries, in total 1446 sequences from publications, PDB entries and GenBank, suggesting it is not regularly updated. The INDI database contains approximately 11 million sequences from nextgeneration sequencing (NGS) studies, and ∼21,000 sequences from patents and literature. The data is available to download, however our analysis (see **Results**) indicates that most of the sequences from patents and literature are false positives, reducing the actual number of VHH sequences from ∼21,000 to below 7000. NanoLAS relies on INDI as a source for their ∼20,000 sequences. Also, to our knowledge, none of these include VNAR data and thus they are missing an important component of the available nanobody data. Here, we present PLAbDab-nano, a database containing 4913 annotated VHH and VNAR sequences from 796 small scale studies. We offer open access to the data and functionality to search the database based on sequence identity using KA-search (*25*) and BLAST (*26*), structural similarity, or keywords. Each sequence is accompanied by a direct link to its source material, facilitating access to further information on any nanobody of interest.

## Methods

### Collecting nanobody sequences

Sequences are drawn from the NCBI GenBank Protein database (*27*), SAbDab (*19, 28*), and TheraSAbDab (*20*) (**Figure 2**). Data is extracted from GenBank by querying the database with keywords, following the methods set out by Abanades *et al*. for the PLAbDab antibody database (*29*). Here we have used nanobody-specific search terms (*nanobody, nanobodies, VHH, VH, shark VNAR, shark novel antigen receptor, single chain antibody, single-chain antibody, single chain antibodies, single-chain antibodies*).

**Figure 2.**
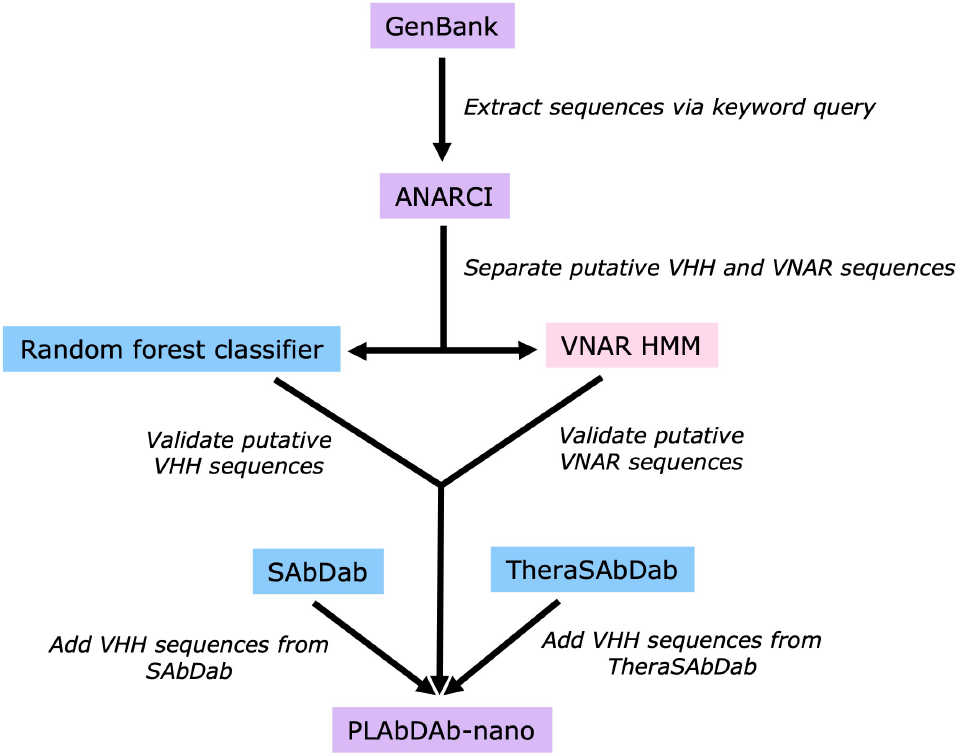
Data is scraped and processed from GenBank (*27*), SAbDab (*19, 30*) and TheraSAbDab (*20*) to generate the PLAbDab-nano database.

Entries resulting from this search are filtered using ANARCI (*30*) to identify VHH sequences, leveraging the Hidden Markov models (HMMs) that support the numbering tool. Since ANARCI was not designed to handle VNAR sequences, separate HMMs were built with a set of manually curated VNAR sequences (sourced from GenBank) to mine VNARs from remaining non-VHH entries. To ensure that VH sequence entries were excluded, we trained a random forest classifier on camel VHH and human and non-human VH sequences, achieving an accuracy of 94%. This model was used to filter out VH sequences.

Further VHH entries were retrieved from SAbDab (*19, 28*) and TheraSAbDab (*20*). TheraSAbDab was filtered for only VHH or single-domain antibody entries, including multispecific therapeutics and at all clinical stages.

### Annotation of VHH and VNAR sequences

ANARCI (*30*) was used to annotate the CDRs of all the VHH sequences. For VNARs, we developed a bespoke annotation method that uses a large language model trained on antibody sequences from multiple species and conditioned on manually curated VNAR sequences. Since VNARs do not have a CDR2, the CDR annotation for VNARs includes only the CDR1 and CDR3 loops. Additional metadata, including antigen information, is added following Abanades *et al*. (*29*), with a small adjustment for species annotation: GenBank species annotation was replaced by that from SAbDab to account for occasions on which non-nanobody chains within an entry caused mislabelling of the origin organism.

### Searching PLAbDab-nano

KA-search (*25*) can be used to carry out a rapid sequence similarity search of VHH entries. Due to the dependency of KA-search on ANARCI, a separate search function for VNARs was built with BLAST (*26*). Users can search over the whole sequence, the CDRs or the CDR3, with or without matching the length of the query and a user-defined sequence identity threshold can be set.

The VHHs were modelled using NanoBodyBuilder2 (*31*), allowing users to search for similar structures. Sequences for which NanoBodyBuilder2 was not able to generate a model, including all VNAR sequences are labelled as ‘FAILED’ in the ‘model’ column of the database. Input query sequences are modelled without refinement using NanoBodyBuilder2 and the framework is aligned to entries with equivalent CDR loop lengths. As in Abanades *et al*., entries are ranked using the carbon-alpha (Cα) root-mean squared deviation (RMSD) over all CDR residues. Lastly, a text search is available for users to search the metadata by keyword.

## Results

### Database statistics

The cumulative number of publicly available sequences has increased substantially over the past decade (**Figure 3**). Currently, the total number of entries in PLAbDab-nano is 4913. Compared to the INDI database (*23*), PLAbDab-nano has fewer entries overall. However, our methods incorporate more stringent filtering. We use a random forest classifier to distinguish VHH from VH sequences, removing the latter from our database. The threshold of 0.55 was calculated using Youden’s J statistic, where 0 corresponds to a VH sequence, and 1 a VHH sequence. To generate our database, we relaxed the threshold to 0.45, to minimise loss of true nanobodies that share a high sequence similarity with VH sequences, and then manually removed just 30 false positive sequences that remained. Applying our model to INDI’s sequences derived from the PDB, GenBank, patents and literature, suggests that, at the 0.55 threshold, almost 70% of their data may be antibody VH sequences and are erroneously included in their output (**SI Figure 1, SI Figure 2**).

**Figure 3.**
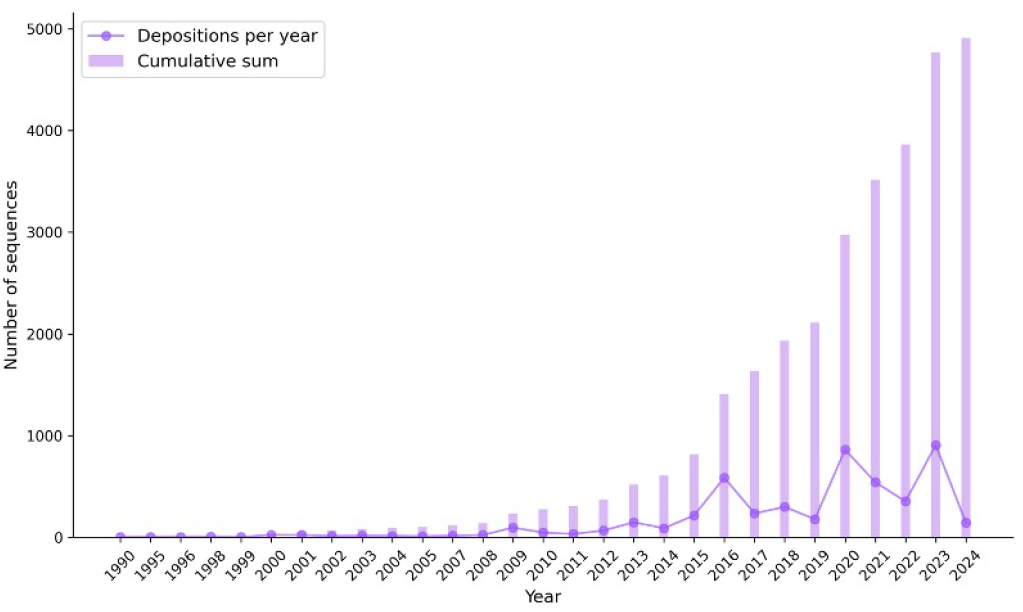
There is an increasing number of depositions of nanobody sequences per year to publicly available sources.

When accounting for redundant sequences, entries in PLAbDab-nano are sourced relatively evenly from the different sources: patents, literature and crystal structures (**Figure 4A**). Excluding patents, due to a lack of species annotation, for VHHs most entries are labelled as llama, followed by alpaca and camel species (**Figure 4B**). For VNARs, entries are split between various shark species (**Figure 4C**). For both VHH and VNAR, approximately 10% of entries come from synthetic constructs.

**Figure 4.**
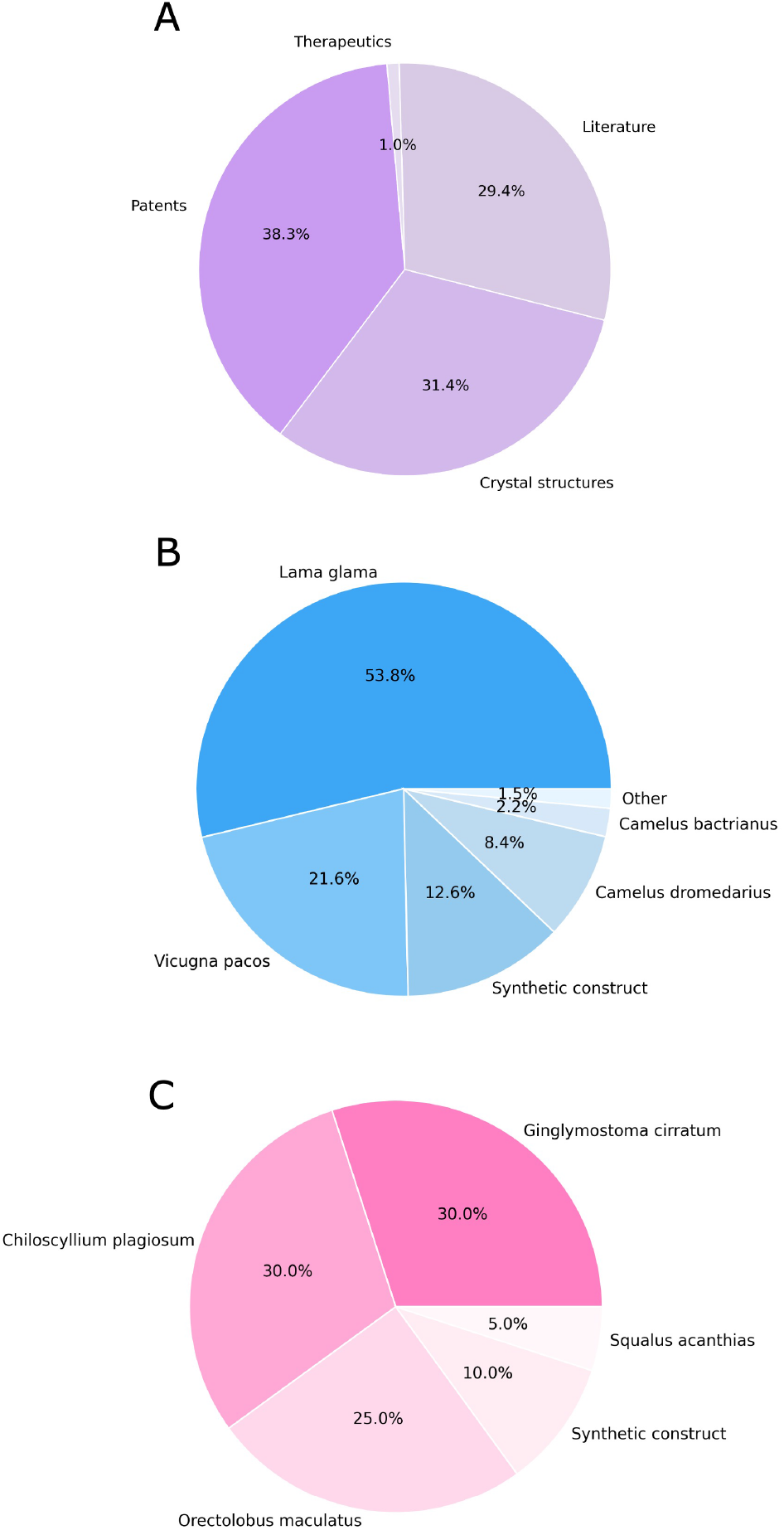
(**A**) Entries in PLAbDab-nano are sourced relatively evenly from the different sources: patents, literature and crystal structures, with a minority from therapeutic data. (**B**) Excluding patents due to a lack of species annotation, most VHH entries are sourced from camelid species and synthetic constructs. (**C**) For VNARs, entries are split between various shark species and synthetic constructs.

The distribution of CDR3 loop lengths from VHH entries in PLAbDab-nano reflects those found in natural immune repertoires (**Figure 5A**), contrasting with the trend observed for natural versus engineered conventional antibodies, where the CDR-H3 in engineered antibodies tend to be shorter (*29*). The CDR3 loops of VNARs tend to be longer than VHH CDR3 loops (**Figure 5B**).

**Figure 5.**
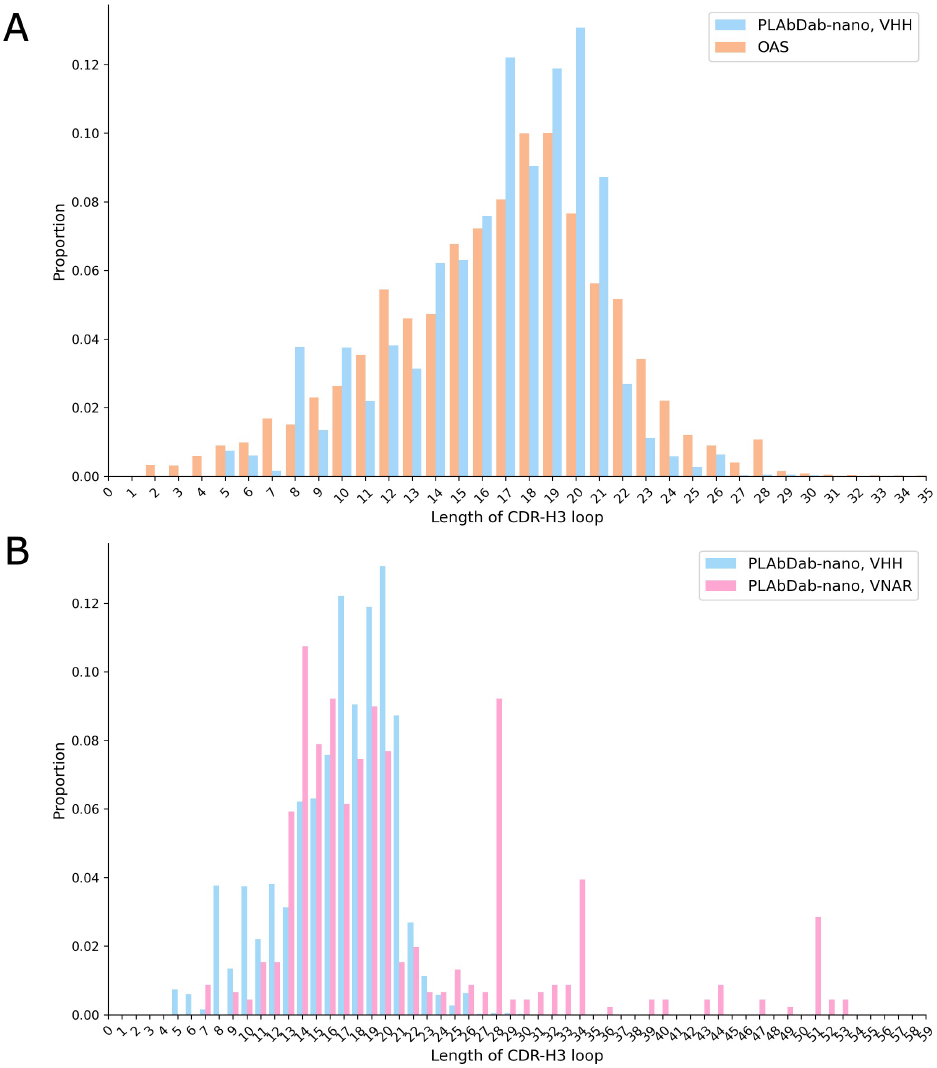
(**A**) The distribution of CDR3 loop lengths (by number of residues) from VHH entries in PLAbDab-nano closely matches those from natural VHH immune repertoire data taken from OAS (*17, 18*) (**B**) For entries included in PLAbDab-nano, the CDR3 loops of VNARs tend to be longer than VHH CDR3 loops.

### VNAR data in PLAbDab-nano provides CDR3 diversity

It has been established in several studies that VHHs have longer CDR3 loops than those occurring in the VH chains of conventional antibodies (*7–12*). VNARs tend to exhibit even longer CDR3 loops than VHHs, further increasing the possible variation in the CDR3, which plays a significant role in binding. Here, we investigated the diversity of the CDR3 loops recorded in the PLAbDab-nano database.

A pairwise comparison of our VHH and VNAR CDR3 loops, including only those that matched by length (34,721 out of 527,058 possible combinations of non-redundant VHH/VNAR pairings) demonstrated low sequence identity between these two types of nanobody (**SI Figure 3**). This would indicate that the majority of VNAR CDR3 loops occupy a distinct sequence space compared to VHHs, and thus nanobodies from sharks provide additional diversity that could be exploited for therapeutic use.

We further assessed the CDR3 diversity from a structural perspective. Using our keyword search and querying for ‘SARS-CoV-2’ in the database, we found 278 entries, 2 of which were VNARs and the remaining 276 VHHs, 1 of which was a therapeutic entry, Rimteravimab. Of the structures available, we found two pairs of VHHs and VNARs that bind at overlapping epitopes on the receptor-binding domain spike protein with very different CDR3 loops (**Figure 6**). These structures all show the CDR3 as dominant in binding their target, but with differing orientations of the scaffold between the two epitopes. This illustrates that the added sequence diversity may also be beneficial in the design of bi- or multi-specific therapeutics where multiple epitopes need to be accessed, as well as, for example, overcoming developability issues such as sequence liabilities.

**Figure 6.**
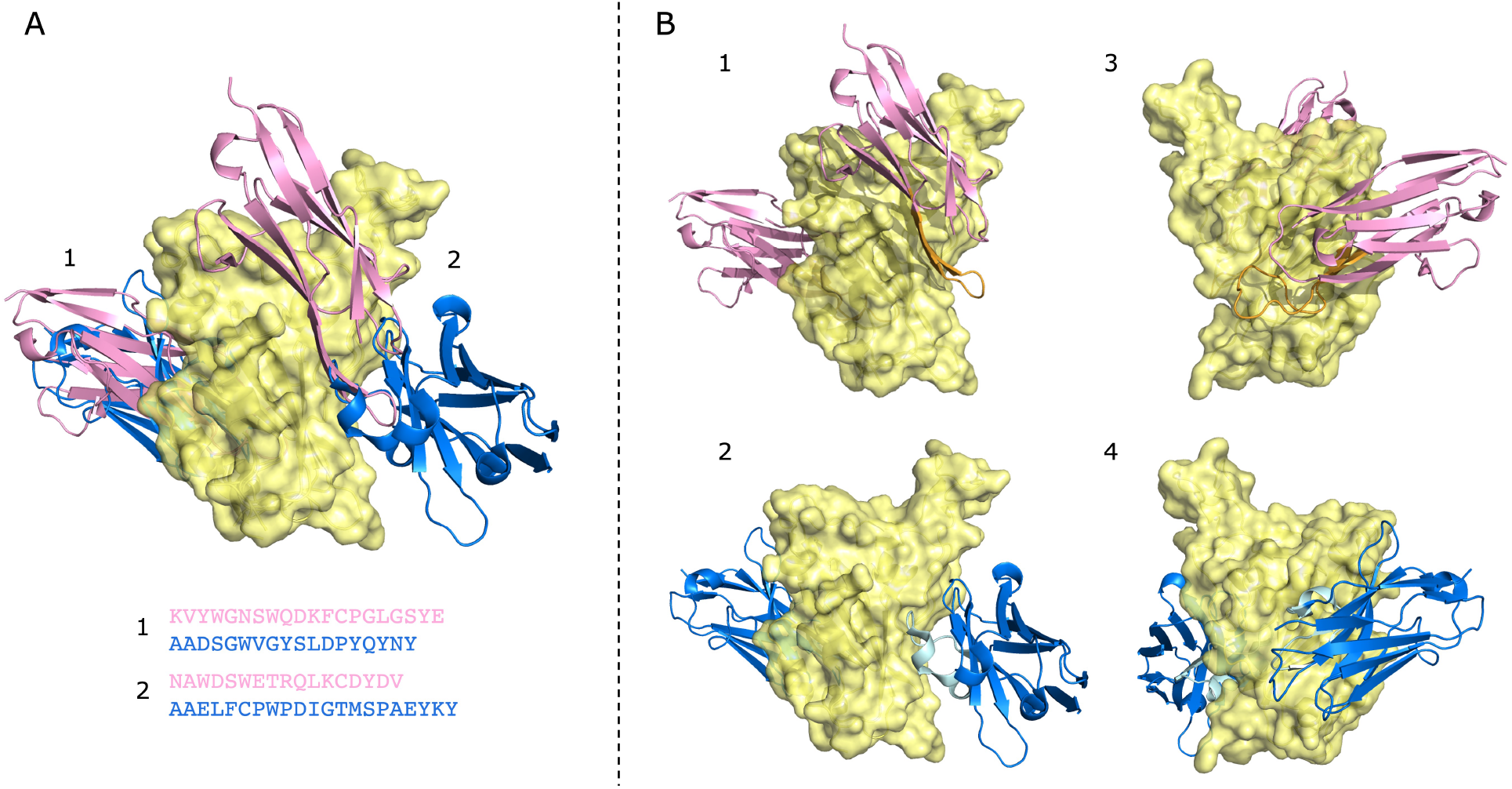
(**A**) VNARs (pink) and VHHs (blue) binding to overlapping epitopes on the RBD of SARS-CoV-2 (yellow). Their CDR3 sequences for two different binding sites are labelled 1 and 2. (**B**) These nanobodies bind to their epitopes in differing orientations, with the CDR3 (shown in orange for the VNAR, and light blue for the VHH) dominant in binding in all cases. Structures 1 and 2 are rotated 180°across the y-axis to produce structures 3 and 4. Data comes from PDB entries 7S83 (for the VNARs) and 8HR2 (for the VHHs), with the RBD from both used to align the structures.

## Discussion

We present PLAbDab-nano, a database of nanobody sequences from patents and literature, including both VHHs and VNARs. PLAbDab-nano is the first database, to our knowledge, to collate VNAR data into one central repository.

PLAbDab-nano can be automatically updated as new publications and patents arise to maintain an up-to-date record of nanobody data. We demonstrate the utility of PLAbDab-nano in providing a record of CDR diversity amongst nanobodies and expect that this could be most useful in designing nanobody libraries. Moreover, collating VNAR data in particular will drive forward our understanding of this nanobody variety and their unique characteristics.

The development of PLAbDab-nano involved the design of a numbering tool for VNARs to enable the annotation of their CDR loops, for which no tool currently exists. Although the strategy described in this paper draws out CDR loops that look reasonable for the structures available, there will be exceptions given the diversity of these regions, particularly for the CDR3 loop. In addition, as was the case for PLAbDab, the resulting database generated by PLAbDab-nano relies on data being publicly available and in a format amenable to web-scraping. As such, since not all sequences are submitted to repositories such as the NCBI or PDB, the volume of data able to be collated is limited where sequences are deposited within the text of papers, or as images.

Despite this, PLAbDab-nano contains just under 5000 annotated nanobody sequences, and will grow with future updates, providing an invaluable resource to the nanobody research community, particularly for studies on VNARs. PLAbDab-nano can be used to generate diverse libraries with properties based on patented and therapeutic nanobodies, and as a dataset to facilitate a greater depth of knowledge into nanobody properties to propel the development of novel therapeutics.

## Data availability

PLAbDab-nano is freely available to download and query at *opig*.*stats*.*ox*.*ac*.*uk/webapps/plabdab-nano/*.

## Acknowledgements

We extend thanks to Brennan Abanades, Tobias Olsen, and the co-authors of the original PLAbDab database (*29*).

