## Supplementary Material for "PLAbDab-nano: a database of camelid and shark nanobodies from patents and literature"

### Supplementary Information

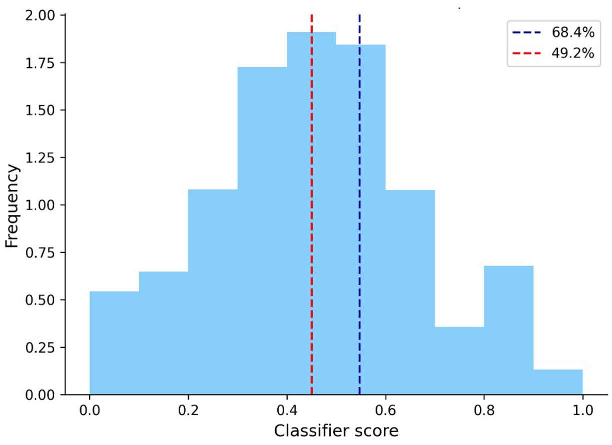


**Figure S1.** The distribution of classifier scores for sequences from patents and literature in the INDI database indicates that 68.4% of entries at our standard threshold of 0.55 (blue dashed line) and 49.2% at a relaxed threshold of 0.45 (red dashed line) would not pass our filtering methods, as they are classified as VH sequences.

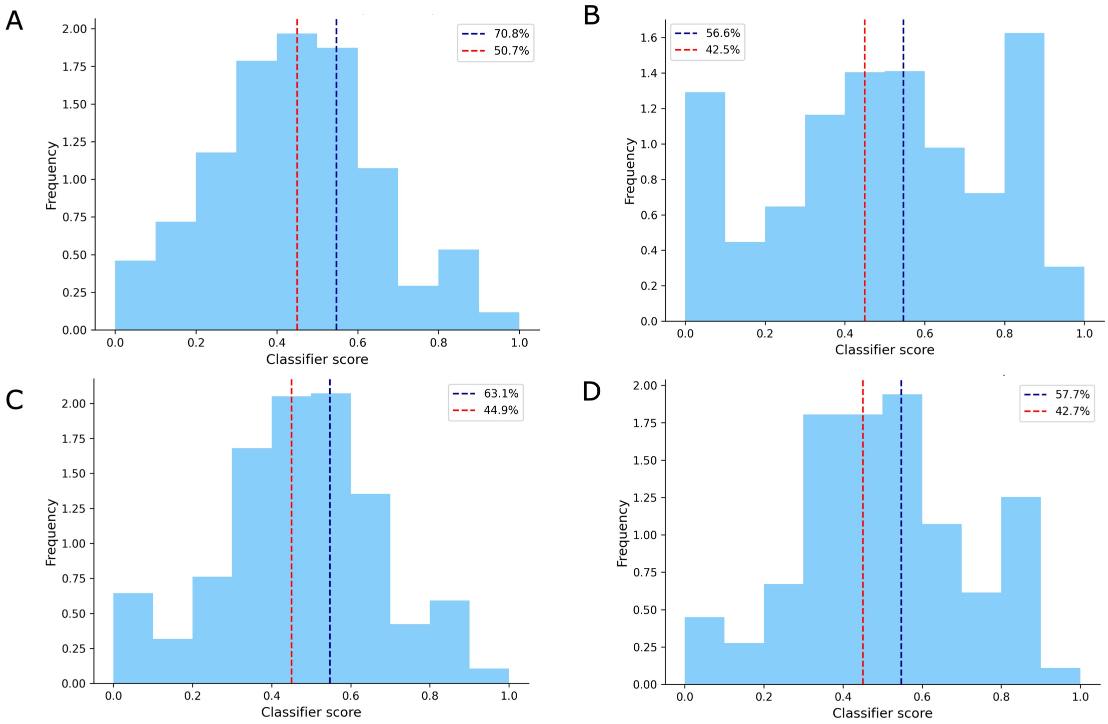


**Figure S2.** Distributions of classifier scores from INDI sequences, split by source into data from (**A**) patents, (**B**) GenBank, (**C**) structures, and (**D**) literature. Entries with scores to the left of the blue dashed lines (threshold = 0.55, calculated using Youden's J-statistic) and red dashed lines (where the threshold was relaxed to 0.45 to account for high VH and VHH sequence similarity) would not pass the filtering methods used to generate PLAbDab-nano, but are included in the INDI database. The values given in the figure legends next to the dashed lines indicate the percentage of sequences that would fail at that threshold.

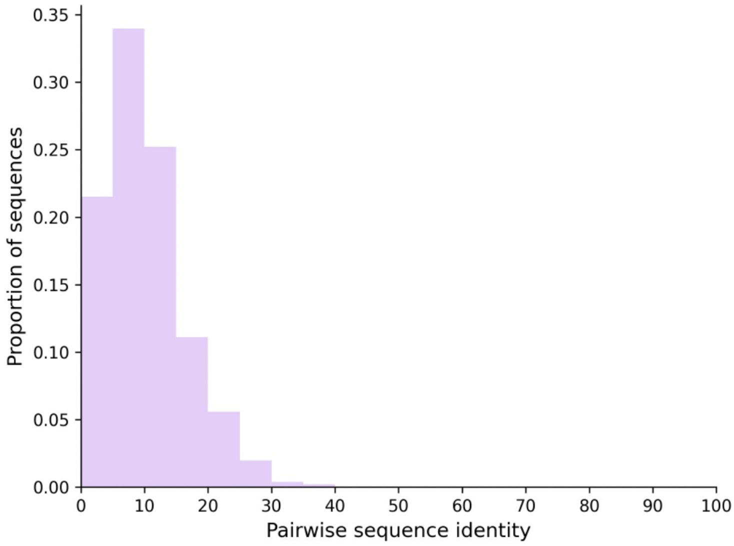


**Figure S3.** Low pairwise sequence identities (by percentage) for VHH CDR3 loops versus VNAR CDR3 loops indicate that the two types of nanobody occupy different regions of sequence space.
